# CD300lf is the primary physiologic receptor of murine norovirus but not human norovirus

**DOI:** 10.1101/859025

**Authors:** Vincent R. Graziano, Forrest C. Walker, Elizabeth A. Kennedy, Jin Wei, Khalil Ettayebi, Madison S. Simões, Renata B. Filler, Ebrahim Hassan, Leon L. Hsieh, Abimbola O. Kolawole, Christiane E. Wobus, Lisa C. Lindesmith, Ralph S. Baric, Mary K. Estes, Robert C. Orchard, Megan T. Baldridge, Craig B. Wilen

## Abstract

Murine norovirus (MNoV) is an important model of human norovirus (HNoV) and mucosal virus infection more broadly. Viral receptor utilization is a major determinant of cell tropism, host range, and pathogenesis. The *bona fide* receptor for HNoV is unknown. Recently, we identified CD300lf as a proteinaceous receptor for MNoV. Interestingly, its paralogue CD300ld was also sufficient for MNoV infection *in vitro*. Here we explored whether CD300lf is the sole physiologic receptor *in vivo* and whether HNoV can use a CD300 ortholog as an entry receptor. We report that both CD300ld and CD300lf are sufficient for infection by diverse MNoV strains *in vitro*. We further demonstrate that CD300lf is essential for both oral and parenteral MNoV infection and to elicit anti-MNoV humoral responses *in vivo*. In mice deficient in STAT1 signaling, CD300lf is required for MNoV-induced lethality. However, after high dose intraperitoneal challenge with MNoV in *Cd300lf*^−/−^*Stat1*^−/−^ mice a single amino acid mutation in the MNoV capsid protein emerged. This substitution did not alter receptor utilization *in vitro*. Finally, we demonstrate that human CD300lf (huCD300lf) is not essential for HNoV infection, nor does huCD300lf inhibit binding of HNoV virus-like particles to glycans. Thus, we report huCD300lf is not a receptor for HNoV.

**Author Summary:** Human norovirus is the leading cause of non-bacterial gastroenteritis causing up to 200,000 deaths each year. How human norovirus enters cells is unknown. Because human norovirus is difficult to grow in the laboratory and in small animals, we use mouse or murine norovirus as a model system. We recently discovered that murine norovirus can use the either CD300ld or CD300lf as a receptor *in vitro*. We also showed that CD300lf deficient mice were resistant to oral challenge with a single virus strain. Here we determined that CD300lf is essential for infection of diverse murine norovirus strains in cell lines and in mice with normal immune systems demonstrating it’s the primary physiologic receptor for diverse murine norovirus strains independent of infection route. However, in immunodeficient mice injected with high dose virus directly into the abdominal cavity, we observed a norovirus mutant that enabled CD300lf-independent infection. Finally, we demonstrated that human CD300lf is not the elusive receptor for human norovirus.

## Introduction

Human norovirus (HNoV) is the leading cause of infectious gastroenteritis globally, yet our understanding of how HNoV enters cells is limited^1–4^. Viral entry is the first and often rate-limiting step of the viral life cycle and is a major determinant of cell tropism, species range, host genetic susceptibility, and pathogenesis^1, 5^. Entry includes virion attachment to the host cell membrane, receptor engagement, internalization, and genome release into the cell cytoplasm^1, 5^. Both histo-blood group antigens (HBGAs) and bile salts bind HNoV and promote infection^6–8^. The identity of a proteinaceous cellular receptor(s) for HNoV remains unknown^1, 8, 9^.

Murine norovirus (MNoV) represents a surrogate animal model for studies of HNoV infection and pathogenesis^1,10, 11^. In contrast to HNoV, diverse infectious molecular clones exist for MNoV that can be readily propagated both *in vitro* and in mice. MNoV and HNoV share similar genome and capsid architecture, fecal-oral transmission routes, interactions with bile salts, and the ability to cause both acute and persistent infection^1, 7, 8, 10–14^. Recently, we identified CD300 family members CD300ld and CD300lf as functional receptors for MNoV^15, 16^. CD300lf is necessary for infection of MNoV-susceptible RAW 264.7 and BV2 cell lines while both CD300ld and CD300lf are sufficient to confer susceptibility to cell lines from different species when ectopically expressed^15, 16^. In addition, CD300lf-deficient mice are resistant to fecal-oral transmission of MNoV strain CR6 (MNoV^CR6^)^15^. Whether CD300lf is essential for genetically diverse MNoV strains, parenteral infection routes, or in the setting of immunodeficiency is unclear. Finally, it is unknown whether human CD300lf functions as a receptor for HNoV.

Noroviruses are non-enveloped, positive-sense single-stranded RNA viruses^1^. The viral capsid exhibits T=3 icosahedral symmetry and is comprised of 90 dimers of the major structural protein VP1^17^. VP1 contains a shell (S) domain and a protruding (P) domain that contains a proximal P1 and distal P2 subdomain^18^. P2 is the target for several neutralizing monoclonal antibodies for both HNoV and MNoV and contains the receptor binding site for MNoV^12, 13^. Co-crystal structures and mutagenesis studies have identified that the CC’ and CDR3 loops of CD300lf directly bind to a cleft between the AB’ and DE’ loops of the MNoV VP1 P2 subdomain^13, 15, 19^. Each CD300lf binds one P2 monomer, albeit at relatively low affinity, suggesting that norovirus-receptor interactions are largely driven by avidity^13^. Interestingly, binding of MNoV to CD300lf is promoted by both divalent cations and bile salts^12, 13, 15^.

Diverse MNoV strains have been described with distinct pathogenic properties^20–22^. Specifically, MNoV^CW3^ causes acute systemic infection that is cleared by the adaptive immune system, while there are other strains including MNoV^CR6^ that cause chronic enteric infection that can last for months if not the life of the animal^20, 23^. Interestingly, a single amino acid in the P2 domain of MNoV VP1 is sufficient to confer lethality in mice deficient in type I interferon signaling, suggesting a role for virus-receptor interactions in determining pathogenesis^20, 24^.

There are eight CD300 family members in mice and seven in humans, each containing a conserved ectodomain, a single transmembrane domain, and a more variable cytoplasmic signaling domain^25, 26^. The ectodomains bind diverse phospholipids found on dead and dying cells, resulting in activating or inhibitory signals depending upon the cellular and molecular context^26–30^. Both CD300ld and CD300lf are expressed in diverse myeloid cells, while CD300lf is additionally expressed in lymphoid cells and rare intestinal epithelial cells called tuft cells, which are the primary target cell of MNoV^CR6^ ^25, 31^.

Here we show that CD300lf is essential for fecal-oral transmission and pathogenesis of diverse MNoV strains in both immunocompetent and immunodeficient mice, suggesting CD300lf is the primary physiologic receptor for MNoV. We also identify CD300lf-independent viral infection in the setting of extra-intestinal challenge in immunodeficient mice, suggesting a potential role for CD300ld or other receptors in this non-physiological infection route. Finally, we demonstrate that CD300lf and related CD300 family members do not function as HNoV receptors.

## Results

HNoV induces vomiting and diarrhea within approximately 24 hours after infection^32–34^. In contrast, MNoV replicates to high titers but is largely avirulent in immunocompetent mice ^10, 35, 36^. Diverse strains of MNoV have been described that can cause either acute self-limiting infection (e.g. MNoV^CW3^) or chronic infection (e.g. MNoV^CR6^) in wild type mice^20, 21^. We previously demonstrated that *Cd300lf*^−/−^ mice do not shed detectable MNoV^CR6^ between 3 and 21 days post-oral challenge; however, the role of CD300lf during earlier time points and in systemic tissues is unclear^15^. MNoV^CW3^ is an infectious molecular clone derived from the MNV-1 plaque isolate CW3^14, 20^. MNoV^CW3^ can infect myeloid and lymphoid cells in the intestine and secondary lymphoid organs and can cause lethal infection in mice deficient in type I interferon signaling^14, 20, 37, 38^. To test whether CD300lf was essential for MNoV^CW3^ infection *ex vivo*, we challenged bone marrow-derived macrophages (BMDMs) from *Cd300lf*^+/−^ or *Cd300lf*^−/−^ mice with MNoV^CW3^. *Cd300lf*^−/−^ BMDMs did not produce infectious virus as measured by plaque assay (Fig 1A) nor did they express the MNoV non-structural protein NS1/2 as detected by flow cytometry (Fig 1B-C). CD300lf is thus essential for MNoV^CW3^ infection of murine BMDMs. To test the role of CD300lf *in vivo*, we challenged *Cd300lf*^+/−^ and *Cd300lf*^−/−^ littermate mice with 10^6^ plaque forming units (PFU) of MNoV^CW3^ perorally (PO) and measured infectious virus by plaque assay (Fig 1D) and viral genomes by qPCR (Fig 1E). At 24 hours post-infection (hpi), infectious virions were undetectable in *Cd300lf*^−/−^ mice in the mesenteric lymph node (MLN), ileum, and colon, in contrast to *Cd300lf*^+/−^ littermates. At this dose and time point, MNoV^CW3^ was not detected in the spleen of control or knockout mice. Viral genomes were similarly undetectable in *Cd300lf*-deficient animals (Fig 1E).

**Figure 1.**
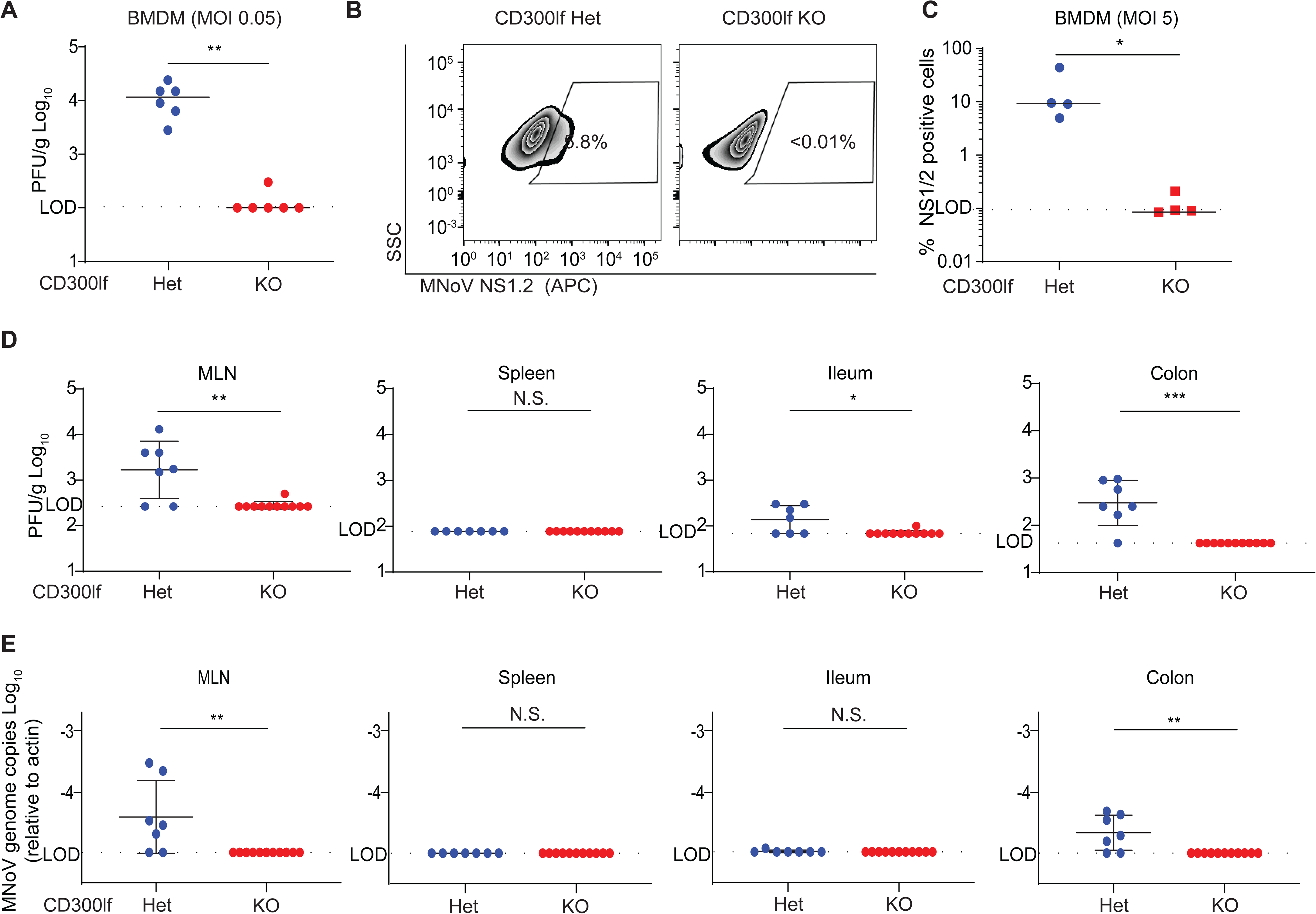
CD300lf is necessary for MNoV^CW3^ infection *ex vivo* and *in vivo*. (A-C) BMDMs were generated from *Cd300lf^−/−^* and *Cd300lf^+/−^* littermate controls. (A) BMDMs were challenged with MNoV^CW3^ (MOI= 0.05) and viral replication was measured by plaque assay 24 hpi. (B) *Cd300lf^+/−^* and *Cd300lf^−/−^* BMDMs were challenged with MNoV^CW3^ (MOI=5) and expression of the MNoV non-structural protein NS1/2 was measured by flow cytometry. (C) Quanitifcation of NS1/2 expression. (D-E) *Cd300lf^+/−^* and *Cd300lf^−/−^* littermates were challenged with 10^6^ PFU PO MNoV^CW3^. 24 hpi, virus was measured by both (D) plaque assay and (E) qPCR in the MLN, spleen, distal ileum, and proximal colon. Data was analyzed by Mann-Whitney test. Shown are means ± SEM. NS, not significant; *P<0.05; **P<0.01; ***P<0.001; L.O.D., limit of detection. Experiments in (A-C) where performed at least two independent times each in triplicate. Data in (D-E) are pooled from two independent experiments with at least three mice per group.

Next, we asked whether CD300lf was required for infection by genetically diverse MNoV strains. MNoV strains are derived from a single genogroup (Genogroup V) and alignments of MNoV VP1 previously revealed at least five distinct clusters, from which we selected representative strains MNoV^CW3^, MNoV^CR6^, MNoV^WU23^, MNoV^CR3^, and MNoV^CR7^ (Table 1) ^21, 39–41^. An alignment of strains highlights the broad amino acid conservation of the capsid protein, including the CD300lf binding sites, and bile-acid interacting sites described previously (Fig S1)^13^. Among these strains, there is no variability in the 11 VP1 residues shown to interact with the secondary bile salts glycochenodeoxycholic acid (GCDCA) or lithocholic acid (LCA)^13^. However, there is variation in 4 of the 17 VP1 residues previously shown to interact with CD300lf, suggesting diverse strains may utilize receptors other than CD300lf (Fig S1)^13^. To test the hypothesis that diverse MNoV strains differentially utilize CD300lf, we first tested whether CD300lf was necessary and sufficient for infection by these strains *in vitro*. We challenged CD300lf-deficient BV2 microglial cells with six diverse MNoV strains. All strains induced cell death in wild-type BV2 cells but not CD300lf-deficient cells, demonstrating CD300lf is essential for virus induced-lethality in BV2 cells (Fig 2A). Next, we asked whether CD300ld and CD300lf are sufficient to confer susceptibility to human HeLa cells. Consistent with our prior findings, both CD300ld and CD300lf are sufficient for infection by MNoV^CW3^ as measured by expression of the MNoV non-structural protein NS1/2 by flow cytometry^15^. Similarly, both CD300ld and CD300lf are sufficient for MNoV^WU23^, MNoV^CR3^, MNoV^CR7^, MNoV^MNV3^, and MNoV^S99^ infection (Fig 2B). All tested MNoV strains utilize CD300ld and CD300lf at similar efficiencies when overexpressed in HeLa cells.

**Figure 2.**
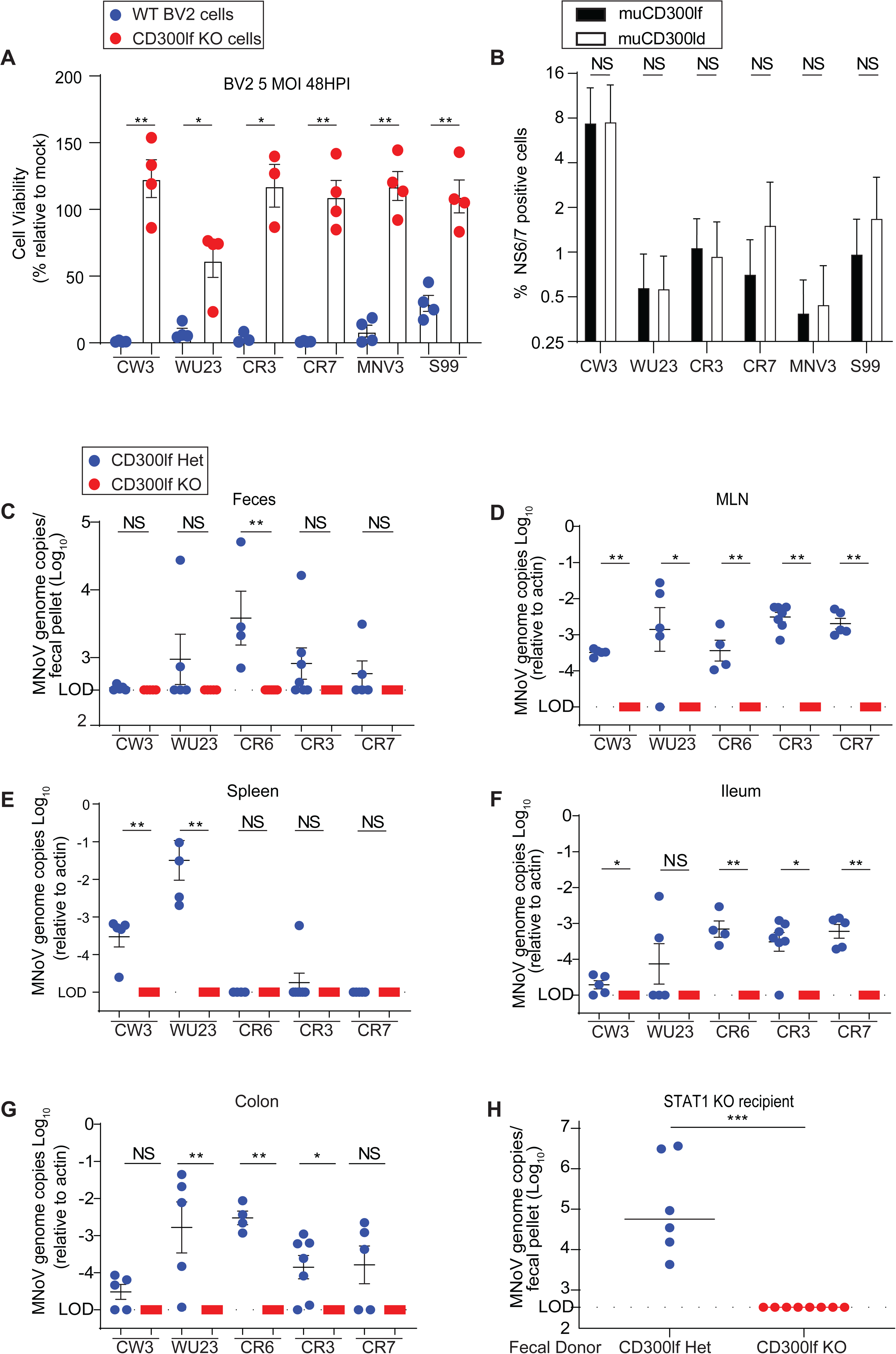
CD300lf is necessary and sufficient for infection by diverse MNoV strains. (A) CD300lf WT and KO BV2 cells were infected with MNoV strains CW3, WU23, CR3, CR7, MNV-3, and S99 at a MOI of 5. CD300lf KO BV2 cells were protected from virus-induced cell death for all MNoV strains. (B) Mouse CD300ld (muCD300ld) and CD300lf (muCD300lf) were overexpressed in human HeLa cells by transient transfection. Cells were challenged with MNoV strains at a MOI of 5 for 24 hours and then infection was quantified by MNoV NS6/7 expression by flow cytometry. (C-H) *Cd300lf^+/−^* or *Cd300lf^−/−^* mice were challenged with 10^6^ PFU PO CW3, WU23, CR6, CR3, or CR7 for seven days. MNoV was detectable in the (C) feces, (D) MLN, (E) spleen, (F) ileum, and (G) colon of *Cd300lf^+/−^* but not *Cd300lf^−/−^* mice. (H) To test whether CR6 was shed in feces below the limit of detection by qPCR, we gavaged *Stat^−/−^* mice with feces from *Cd300lf^+/−^* or *Cd300lf^−/−^* mice challenged with MNoV^CR6^ from (C). Fecal pellets from *Cd300lf^−/−^* mice, did not establish detectable infection in *Stat1^−/−^* mice. Data is pooled from two to four independent experiments. Data was analyzed by Mann-Whitney test. Shown are means ± SEM. NS, not significant; *P<0.05; **P<0.01; ***P<0.001; L.O.D., limit of detection.

**Table 1.**
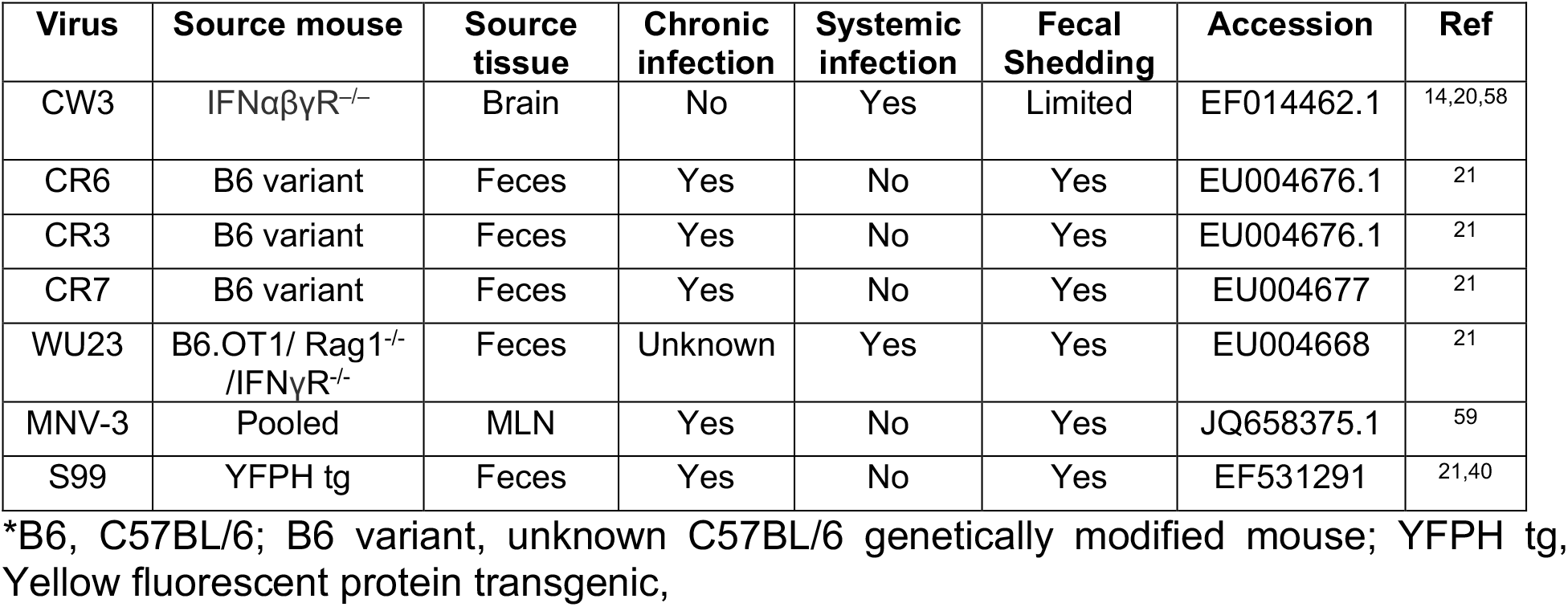

To determine whether CD300lf was essential for infection *in vivo*, we challenged *Cd300lf*^+/−^ and *Cd300lf*^−/−^ littermates with 10^6^ PFU PO of either MNoV^CW3^, MNoV^CR6^, MNoV^WU23^, MNoV^CR3^, or MNoV^CR7^ for seven days. These viruses represent each of the five distinct MNoV VP1 phylogenetic clusters described previously^39^. MNoV^CW3^ causes acute systemic infection that is cleared by 10-14 days post-infection (dpi), while MNoV^CR6^, MNoV^CR3^, and MNoV^CR7^ cause persistent infection^21^. The *in vivo* characteristics of MNoV^WU23^ are uncharacterized^21^. Consistent with prior studies, we observed differential tissue tropism associated with different MNoV strains^21^. In control mice, viral genomes for all tested MNoV strains were shed in the feces of some mice with the exception of MNoV^CW3^ (Fig 2C). All tested MNoV strains were detected in the MLN (Fig 2D). All strains were detectable albeit at varying levels in the ileum and colon. Interestingly, only MNoV strains MNoV^CW3^ and MNoV^WU^^23^ were detectable in the spleen. MNoV^CR6^, MNoV^CR3^, and MNoV^CR7^, which are known to establish chronic infection in immunocompetent mice did not have detectable viral genomes in the spleen. MNoV genomes were not detected in any tissue (MLN, spleen, ileum, colon, or feces) of any *Cd300lf*-deficient mice challenged with any of the five MNoV strains (Fig 2C-G). To determine whether there was MNoV infection below the limit of detection, we collected feces from *Cd300lf*^+/−^ or *Cd300lf*^−/−^ mice challenged with MNoV^CR6^ seven days prior, and then gavaged the feces into *Stat1*^−/−^ recipient mice which lack innate immunity and are exquisitely sensitive to MNoV infection^14, 15^. Seven days after fecal gavage we detected MNoV^CR6^ in *Stat1*^−/−^ mice challenged with feces from *Cd300lf*^+/−^ mice but not from *Cd300lf*^−/−^ mice (Fig 2H). These results suggest CD300lf is critical for infection by multiple MNoV strains in immunocompetent mice *in vivo*.

To further determine whether CD300ld or an alternative receptor was sufficient for MNoV infection below our limit of detection, we assessed the humoral immune response against MNoV^CW3^ at 14 days dpi, a time point after viral clearance by the adaptive immune system. A 1:10 dilution of sera from *Cd300lf*^+/−^ mice that cleared MNoV^CW3^ infection neutralized MNoV^CW3^ *in vitro* as measured by BV2 cell viability (Fig 3A). The IC_50_ from *Cd300lf*^+/−^ mice was approximately 1:80 in contrast to sera from *Cd300lf*^−/−^ mice which did not reach an IC_50_ (Fig 3B). To determine whether MNoV^CW3^ elicited a non-neutralizing humoral response in *Cd300lf*^−/−^ mice, we measured anti-MNoV IgG and IgM in the sera. *Cd300lf*^+/−^ mice had a significantly higher anti-MNoV IgG and IgM response consistent with resistance of *Cd300lf*^−/−^ mice to MNoV^CW3^ infection (Fig 3C-D).

**Figure 3.**
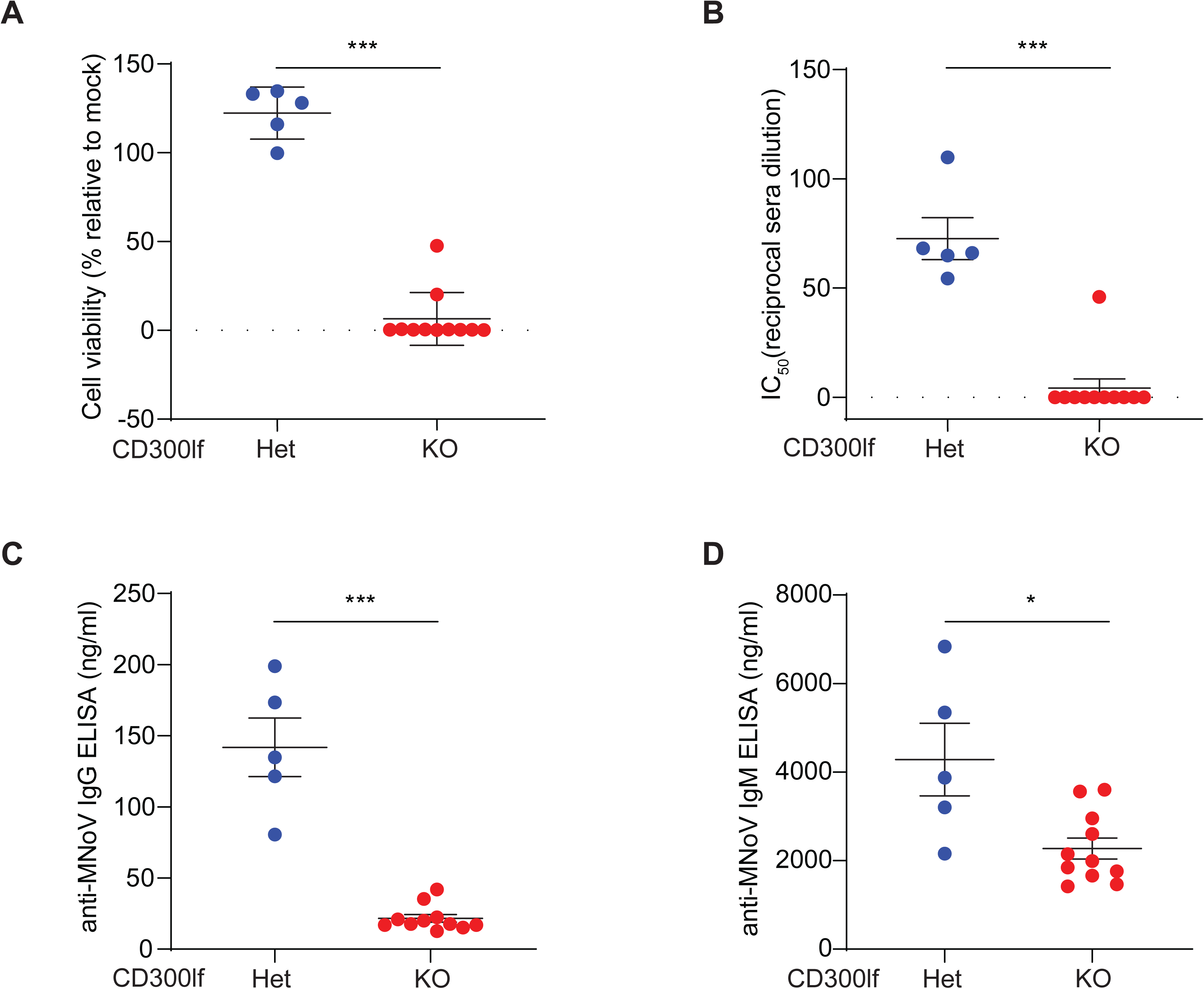
CD300lf is required to generate humoral response after oral MNoV^CW3^ challenge. Sera was collected from *Cd300lf^+/−^* and *Cd300lf^−/−^* mice 14 days after challenge with 10^6^ PFU PO MNoV^CW3^. (A) The maximal protection (1:10 sera dilution) and (B) sera IC_50_ was measured by *in vitro* MNoV^CW3^ neutralization assay in BV2 cells. (C) *Cd300lf^+/−^* generated significantly increased anti-MNoV IgG (C) and IgM (D). Data is pooled from two independent experiments. Data was analyzed by Mann-Whitney test. Shown are means ± SEM. NS, not significant; *P<0.05; **P<0.01; ***P<0.001; ****P<0.0001. L.O.D., limit of detection.

Next, we asked whether CD300lf was essential for fecal-oral transmission of MNoV^CW3^ in *Stat1*^−/−^ mice. We challenged *Cd300lf*^+/+^, *Cd300lf*^+/−^ or *Cd300lf*^−/−^ mice on a *Stat1*^−/−^ background with 10^6^ PFU MNoV^CW3^ PO, which is greater than 1000-fold above the lethal dose^15^. All *Cd300lf*^+/+^*Stat1*^−/−^ succumbed to lethal infection by five dpi (Fig 4A). *Cd300lf*^+/−^*Stat1*^−/−^ similarly all died but with a one-day delay in death suggesting a modest CD300lf gene dosage effect on MNoV pathogenesis. In contrast, all ten *Cd300lf*^−/−^*Stat1*^−/−^ mice survived until at least 21 dpi without overt clinical manifestations (Fig 4A).

**Figure 4:**
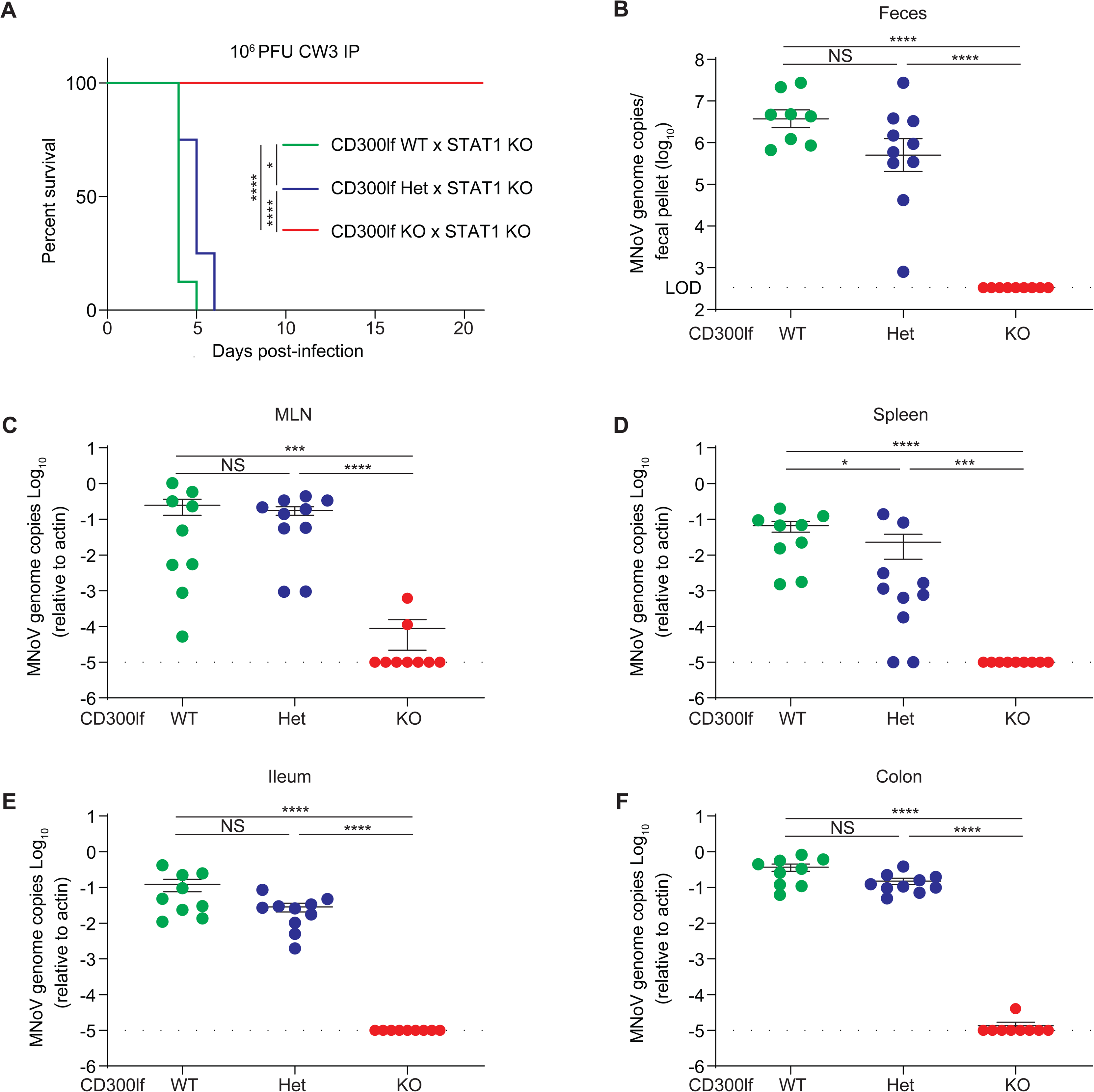
CD300lf is essential for oral MNoV transmission in *Stat^−/−^* mice. **(A)** *Cd300lf^+/+^Stat1^−/−^* (N=8) and *Cd300lf^+/−^Stat1^−/−^* (N=4) mice challenged with 10^6^ PFU PO MNoV^CW3^ succumbed to infection by 5 and 6 dpi, respectively. In contrast, all *Cd300lf^−/−^ Stat1^−/−^* (N=10) mice survived for at least 21 dpi. (B-F) *Cd300lf^+/+^Stat1^−/−^*, *Cd300lf^+/−^stat1^−/−^*, and *Cd300lf^−/−^Stat1^−/−^* mice were challenged with 10^6^ PFU PO MNoV^CR6^. All mice survived infection. Viral genomes were quantified in the (B) feces, (C) MLN, (D) spleen, (E) ileum, and (F) colon at seven dpi. Data is pooled from at least three independent experiments. Data was analyzed by Mann-Whitney test and Kaplan-Meier curves were generated for survival experiments. Shown are means ± SEM. NS, not significant; *P<0.05; **P<0.01; ***P<0.001; ****P<0.0001. L.O.D., limit of detection.

Given the distinct target cell types, tissue tropism, and pathogenesis between MNoV strains, we next tested whether *Cd300lf*^−/−^*Stat1*^−/−^ mice were susceptible to MNoV^CR6^ infection. In contrast to MNoV^CW3^, MNoV^CR6^ is not lethal in *Stat1*^−/−^ mice although the virus replicates to higher titers and can spread to extra-intestinal tissues^14^. All *Cd300lf*^+/+^*Stat1*^−/−^ and *Cd300lf*^+/−^*Stat1*^−/−^ mice challenged with 10^6^ PFU PO MNoV^CR6^ survived the seven-day infection. Viral genomes were detected in the feces, MLN, spleen, ileum, and colon (Fig 4B-F). There was a modest CD300lf gene dosage effect in the spleen (p<0.05) among *Cd300lf*^+/+^ and *Cd300lf*^+/−^ mice, but this was not statistically significant in the MLN, ileum, colon, or feces (Fig 4C-F). Relative to *Stat1*^+/+^ mice, viral genome copies were elevated and extra-intestinal spread to the spleen was observed consistent with prior findings (Fig 2C-G). Importantly, *Cd300lf*^−/−^*Stat1*^−/−^ mice were resistant to MNoV^CR6^ in all tissues examined. Two *Cd300lf^−/−^Stat1^−/−^* mice had detectable viral genomes in the MLN but not in other tissues (Fig 4C). A third mouse had detectable low-level viral RNA in the colon but not in any other tissue, possibly reflective of input virus or a false positive (Fig 4F).

The natural defenses of the gastrointestinal mucosa represent a bottleneck to viral infection. To test whether CD300lf was essential for parenteral infection routes, we bypassed the gastrointestinal tract by administering 10^7^ PFU MNoV^CW3^ intraperitoneally (IP) to *Cd300lf*^−/−^*Stat1*^−/−^ mice. All *Cd300lf*^+/+^*Stat1*^−/−^ and *Cd300lf*^+/−^*Stat1*^−/−^ mice died by five or six dpi, respectively (Fig 5A). Two of nine *Cd300lf*^−/−^*Stat1*^−/−^ mice died at 10 and 14 dpi with the remaining mice surviving at least 21 days. The two *Cd300lf*^−/−^*Stat1*^−/−^ mouse deaths were observed in independent experiments. Virological data was not available from these animals; therefore, whether MNoV^CW3^ contributed to the death of these two *Cd300lf*^−/−^*Stat1*^−/−^ mice is unknown.

**Figure 5.**
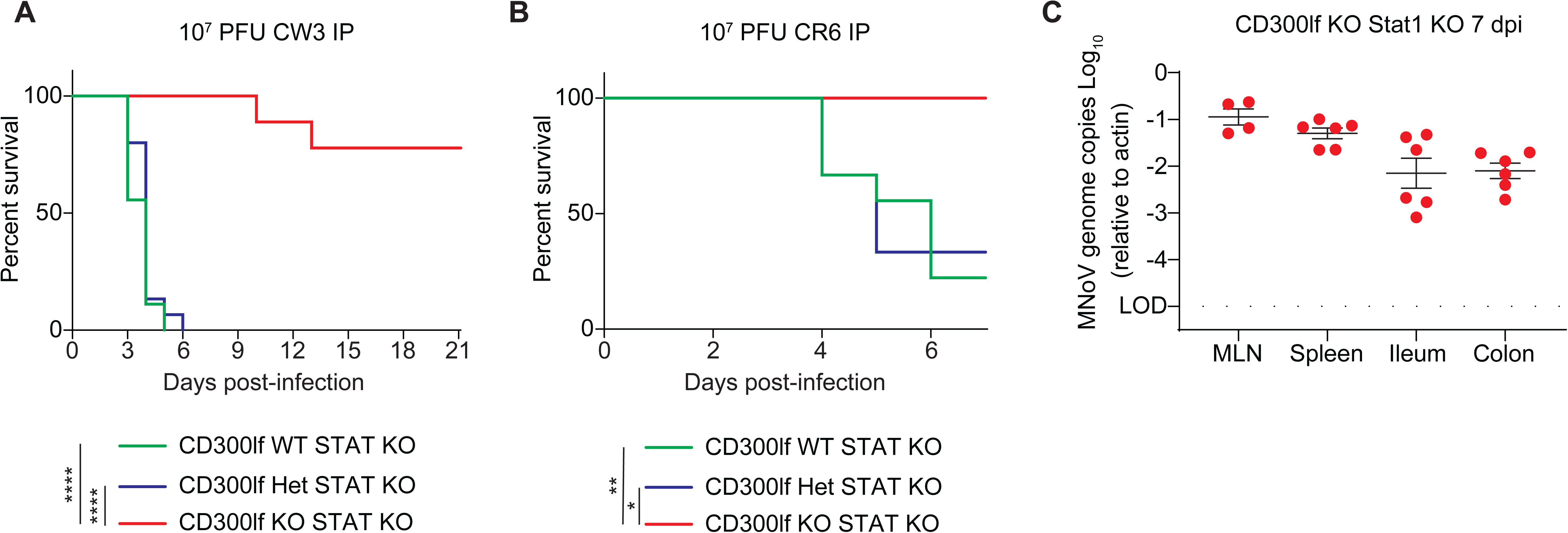
CD300lf is essential for pathogenesis of parenterally transmitted MNoV in STAT1 deficient mice. (A) Mice were challenged with 10^7^ PFU IP MNoV^CW3^. *Cd300lf^−/−^ Stat1^−/−^* mice (N=9 mice) survived infection in contrast to *Cd300lf^+/+^Stat1^−/−^* (N=15 mice) and *Cd300lf^+/−^Stat1^−/−^* (N=9 mice) littermates. (B) Mice were challenged with 10^7^ PFU IP MNoV^CR6^. *Cd300lf^−/−^Stat1^−/−^* mice (N=6 mice) survived infection in contrast to *Cd300lf^+/+^Stat1^−/−^* (N=9 mice) and *Cd300lf^+/−^Stat1^−/−^* (N=3 mice) littermates. (C) MNoV genomes were quantified from the MLN, spleen, ileum, and colon of *Cd300lf^−/−^Stat1^−/−^* mice seven days post-challenge with 10^7^ PFU IP MNoV^CR6^. Data was analyzed by Kaplan-Meier curve for survival experiments. Data is pooled from at least three independent experiments with 1-5 mice per group. *P<0.05; **P<0.01; ***P<0.001; ****P<0.0001. L.O.D., limit of detection.

Next, we challenged mice with 10^7^ PFU MNoV^CR6^ IP. In contrast to the lower-dose MNoV^CR6^ administered PO, 77% (7/9) of *Cd300lf^+/+^Stat1^−/−^* mice and 66% (2/3) *Cd300lf^+/−^Stat1^−/−^* mice died (Fig 5B). All six *Cd300lf*^−/−^*Stat1*^−/−^ mice survived IP infection with 10^7^ MNoV^CR6^ until sacrifice at seven dpi. Interestingly, MNoV^CR6^ genomes were detected in the MLN, spleen, ileum, and colon of *Cd300lf*^−/−^*Stat1*^−/−^ mice (Fig 5C). MNoV^CR6^ from *Cd300lf*^−/−^*Stat1*^−/−^ mice infected with 10^7^ PFU MNoV^CR6^ IP was Sanger sequenced, revealing a single nucleotide change (T6595A) resulting in a phenylalanine to isoleucine mutation at position 514 (F514I) in the P1 domain of VP1 (Supplemental Table 1). This mutation was detected in the spleen, ileum, and colon from two different mice. Thus, while CD300lf is essential for MNoV infection by oral inoculation in immunocompetent hosts, CD300lf-independent replication of MNoV may occur under high-dose IP challenge in *Stat1^−/−^* mice.

To determine whether the F514I variant observed enabled CD300lf-independent infection, we generated an infectious molecular clone containing this mutation on the MNoV^CR6^ background (MNoV^F514I^). Similar to MNoV^CR6^, MNoV^F514I^ was cytotoxic to WT BV2 cells but did not affect cell viability in *Cd300lf*-deficient BV2 cells, demonstrating this mutation did not enable alternative receptor utilization in BV2 cells (Fig 6A). We then asked whether MNoV^F514I^ could preferentially utilize CD300ld relative to CD300lf when ectopically expressed in HeLa cells. MNoV^F514I^ and MNoV^CR6^ were both able to utilize CD300ld and CD300lf at similar ratios when overexpressed suggesting this mutation does not enhance CD300ld utilization *in vitro* (Fig 6B). Next, we thermally stressed MNoV^CR6^, MNoV^F514I^, MNoV^CW3^ to probe differences in virion stability as mutations that confer heat resistance were recently demonstrated to alter the VP1 conformation and thus may affect receptor utilization^42^. MNoV^F514I^ and MNoV^CR6^ had similar thermal stability (Fig 6C). Interestingly, this was increased relative to MNoV^CW3^ (Fig 6C). To investigate whether MNoV^F514I^ altered receptor utilization *in vivo*, we challenged *Cd300lf^+/−^* or *Cd300lf^−/−^* mice with 10^6^ PFU MNoV^F514I^ PO. MNoV^F514I^ readily infected *Cd300lf^+/−^* mice and maintained similar tissue tropism as the parental MNoV^CR6^ (Fig 6D-G). Also similar to MNoV^CR6^, MNoV^F514I^ genomes were not detectable in the MLN, spleen, ileum, colon, or feces of CD300lf^−/−^ mice suggesting MNoV^F514I^ is not sufficient to alter receptor utilization in immunocompetent mice when administered PO (Fig 6D-H).

**Figure 6.**
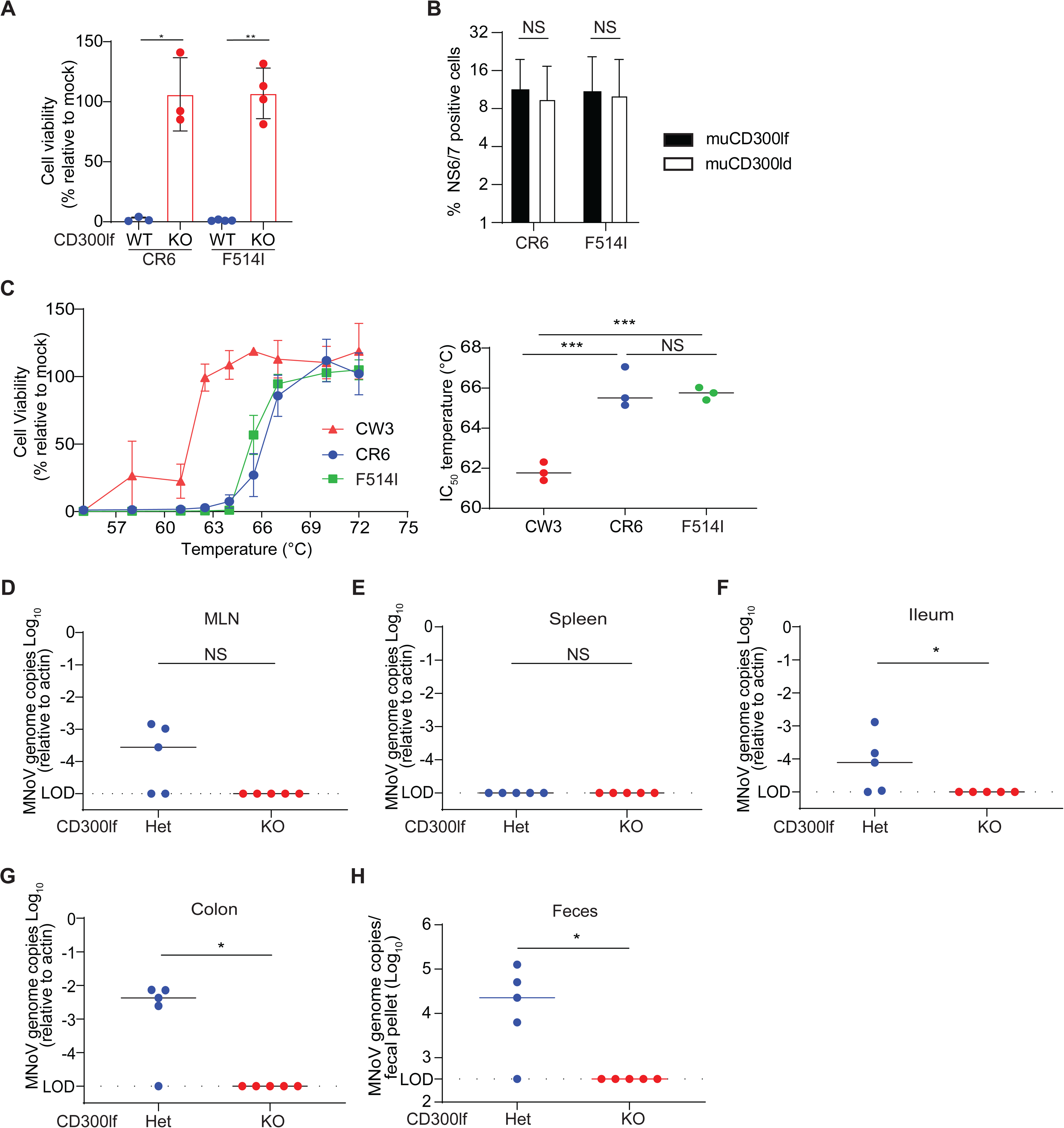
Emergence of MNoV^F514I^ variant after high dose intraperitoneal challenge in *Cd300lf^−/−^Stat1^−/−^* mice. Sanger sequencing revealed a single amino acid mutation in the P1 domain of MNoV^CR6^ after IP challenge with 10^7^ PFU MNoV^CR6^ in *Cd300lf^−/−^Stat1^−/−^* mice. An infectious molecular clone of MNoV^F514I^ was generated. (A) MNoV^F514I^ mediated cell death of BV2 cells is CD300lf dependent. (B) HeLa cells transiently expressing murine CD300ld (muCD300ld) or murine CD300lf (muCD300lf) where challenged with MNoV^CR6^ or MNoV^F514I^ (MOI of 5). MNoV infection was determined by expression of MNoV NS6/7 by flow cytometry. MNoV^F514I^ and MNoV^CR6^ similarly utilize muCD300ld and muCD300lf when overexpressed. (C) MNoV^F514I^ and MNoV^CR6^ have similar thermal stability which differs from MNoV^CW3^. (D) *Cd300lf^+/−^* and *Cd300lf^−/−^* mice were challenged with 10^6^ PFU PO MNoV^F514I^. Viral genomes were quantified in the (D) MLN, (E) spleen, (F) ileum, (G) colon and (H) feces at seven dpi. MNoV^F514I^ infection was detected in *Cd300lf^+/−^* mice but not *Cd300lf^−/−^* mice. Data was analyzed by Mann-Whitney tests. Shown are means ± SEM. NS, not significant; *P<0.05; **P<0.01; ***P<0.001; L.O.D., limit of detection.

Next, given the structural and genetic similarity between MNoV and HNoV, we tested whether human CD300lf (huCD300lf) is a receptor for HNoV. First, we assessed the ability of huCD300lf to prevent binding of recombinant HNoV virus-like particles (VLPs) to HBGAs in pig gastric mucin. To increase the avidity of the potential CD300lf and VLP interaction, we generated Fc-fusion proteins with the either huCD300lf or human CD300ld (huCD300ld) ectodomains. We recently demonstrated that an Fc-fusion protein of mouse CD300lf has substantially increased binding and neutralizing ability on MNoV^13^. We incubated HNoV VLPs with 10 µg/ml of either Fc-huCD300lf, Fc-huCD300ld, or a buffer-only control and measured HBGA-bound VLPs by ELISAs. Neither Fc-huCD300lf or Fc-huCD300ld inhibited binding of GI.1 Norwalk, GI.3 Desert Shield Virus, GII.4 1997, or GII.4 2012 VLPs to HBGAs (Fig 7A)^43, 44^. Next, we asked whether potential HNoV interactions with huCD300lf were necessary for infection of human intestinal enteroids (HIEs). Differentiated monolayers of HIEs were incubated with polyclonal anti-huCD300lf or an IgG1 control and then challenged with HNoV GII.4 from stool filtrate. Anti-huCD300lf had no effect on HNoV genome replication at 24 hpi relative to the isotype control (Fig 7B). Finally, we asked whether Fc-huCD300lf could neutralize HNoV GII.4. Virus was pre-incubated with up to 50 µg/ml of Fc-huCD300lf and then used to infect HIEs. Fc-huCD300lf did not affect viral genome replication relative to a control protein (Fig 7C). Together these data indicate that huCD300lf is not a receptor for HNoV.

**Figure 7.**
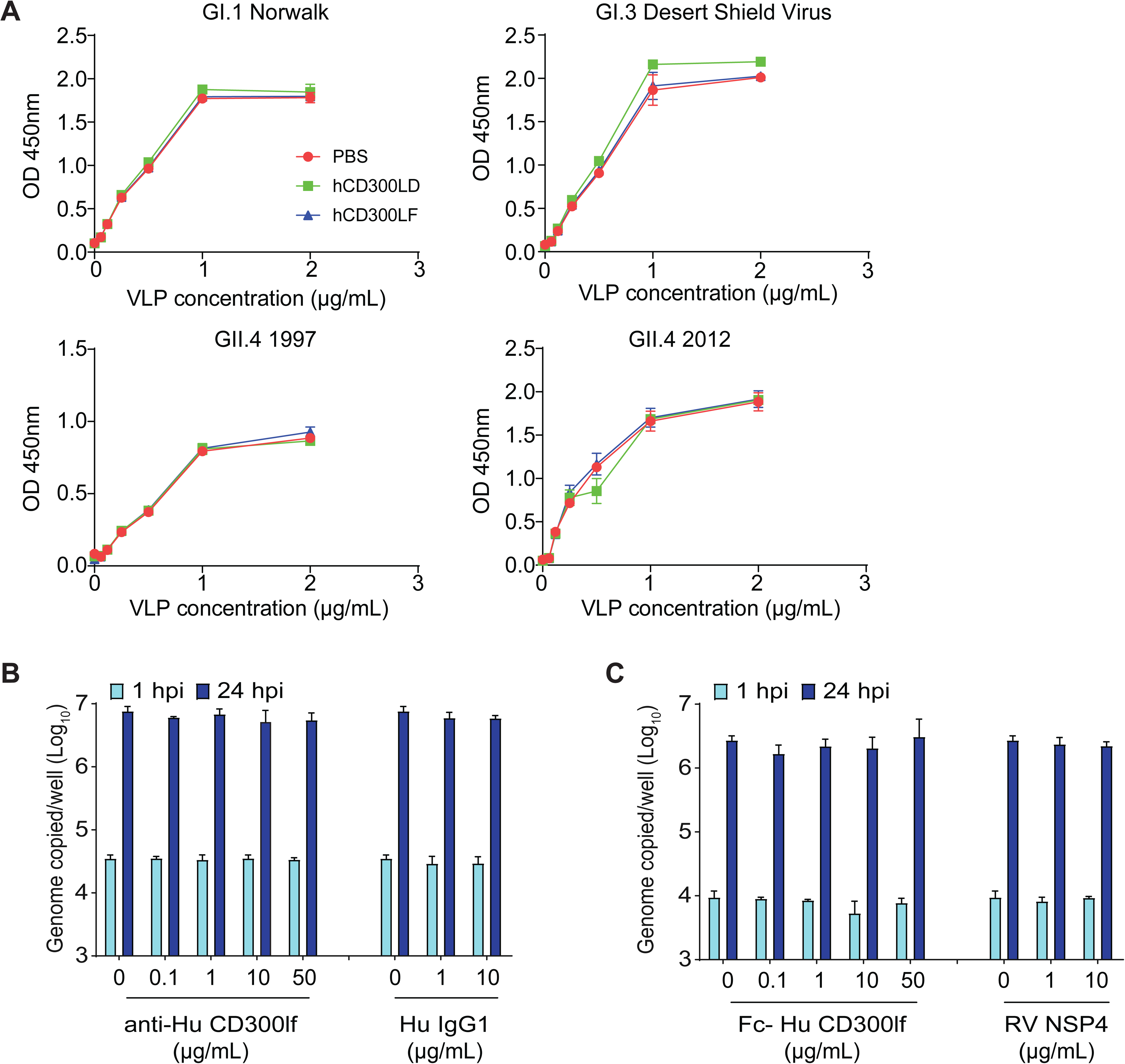
Human CD300lf is not a HNoV entry factor. (A) Human CD300lf and human CD300ld Fc-fusion proteins do not prevent binding of HNoV virus-like particles (VLPs) to pig gastric mucin for GI.1 Norwalk, GI.3 Desert Shield Virus, GII.4.1997, and GII.4.2012. (B) Polyclonal antibody against human CD300lf does not prevent HNoV GII.4 replication in HIEs relative to an IgG1 control. (C) Pre-incubating HNoV GII.4 with a human CD300lf Fc-fusion protein does not prevent HNoV replication in HIEs relative to a control protein (RV NSP4). Data in (A) is representative of at least two independent replicates each performed in duplicate. Data in (B and C) is pooled from three independent experiments with each condition and time point performed in triplicate wells of HIE cultures. Shown are means ± standard deviation.

## Discussion

Here we demonstrate that CD300lf, and not CD300ld, is the primary physiologic receptor for diverse MNoV strains *in vivo*. We demonstrated that CD300lf is essential for detectable viral infection at multiple time points with diverse MNoV strains as measured by plaque assay, qPCR, and serologic response. Interestingly, both CD300ld and CD300lf are sufficient to mediate MNoV entry when ectopically expressed *in vitro* yet CD300ld is not sufficient for oral or parenteral transmission in immunocompetent mice. Several possibilities may explain this discordance. First, CD300ld receptor utilization has only been described when overexpressed at supraphysiologic levels^15, 16^. Similarly the soluble recombinant ectodomain of CD300lf can neutralize MNoV infection in contrast to that of CD300ld^15^. This raises the possibility that CD300ld utilization by MNoV is inefficient relative to CD300lf and that our overexpression assays are not sufficiently sensitive to quantify differences in receptor utilization efficiency. A similar phenomenon has been described with HIV-1, which can engage co-receptors other than CCR5 and CXCR4 when overexpressed in cell lines, but not under physiologically relevant conditions^45, 46^. Second, CD300ld and CD300lf have overlapping yet distinct cell type expression, raising the possibility that CD300ld-expressing cells are either not permissive to MNoV because of a post-entry restriction or because CD300ld-expressing cells are not anatomically accessible to MNoV. Consistent with this, intestinal tuft cells, which are the major target of MNoV^CR6^ in the intestines, express CD300lf but not CD300ld^31, 47^.

Although we demonstrated that *Cd300lf*-deficient mice are resistant to diverse MNoV strains via both oral and parenteral routes, we observed detectable viral genomes in *Cd300lf^−/−^Stat1^−/−^* mice inoculated with high-dose MNoV^CR6^ IP, consistent with CD300lf-independent viral replication. Sanger sequencing of virus from tissues from multiple mice revealed a single amino acid mutation (F514I) in the P1 domain of MNoV VP1. Surprisingly, CD300lf remained essential for MNoV^F514I^ infection both *in vitro* and in immunocompetent mice, raising the question as to why this viral variant emerged and whether it has physiologic relevance. One possibility is that MNoV^F514I^ enables the virus to better utilize CD300ld, or an unknown alternative receptor *in vivo*, but only in a *Stat1-* deficient background. Defects in innate immunity may enable broader cell tropism due to removal of post-entry viral restriction, thus permitting CD300lf-independent replication *in vivo*. The molecular consequences of F514I are intriguing in that this mutation is on the opposite side of the P domain from the receptor binding site, suggesting an allosteric mechanism of receptor regulation. One such mechanism could be altering the “breathability” of the P domain, which is connected to the capsid shell by a flexible linker^12, 48^. Bile salts both increase MNoV-receptor binding and induce collapse of the P domain onto the shell^12^. Whether F514I has a similar effect remains an important future direction.

The physiologic function of CD300lf is as a cell death sensor and immunoregulatory protein^25^. Interestingly, the CD300 locus is under positive selection and there are a number of CD300lf polymorphisms in laboratory and wild mouse strains that may affect CD300lf expression or conformation and thus may regulate susceptibility to MNoV^49^. This raises the intriguing hypothesis of a molecular arms race between MNoV and mice over evolutionary time which may have contributed to the relative avirulence of MNoV in immunocompetent mice.

Finally, these data have several important implications for our understanding of HNoV, the entry mechanisms of which remain to be defined. Given the structural similarity between MNoV and HNoV as well as mouse and human CD300lf, we hypothesized that human CD300lf might be a functional receptor for HNoV. However, via multiple orthogonal approaches we demonstrated that human CD300lf is not a functional receptor for several GI and GII HNoVs. This further raises the question as to the identity of the HNoV receptor and whether or not it is structurally related to CD300lf. This remains an important area of future investigation that would significantly enhance our understanding of HNoV pathogenesis and facilitate novel prophylactic and therapeutic approaches.

## Materials and Methods

### Mouse strains

All mouse strains used in this experiment were from a C57BL/6J background (Jackson Laboratories, Bar Harbor, ME) and bred in-house. Generation of B6.CD300lf^em1Cbwi^/J (*Cd300lf^−/−^*) mice (Jackson Laboratories) and B6.129S(Cg)-Stat1^tm1Div^/J (*Stat1^−/−^*) (gift of H.W. Virgin) were previously described^15, 50^. These mice were housed in a MNoV-free facility at Yale University School of Medicine. All experiments used littermate controls, were gender-balanced, and done at least two independent times. Mice were used for infections between 6-10 weeks of age. Genotyping of mice was done by real time PCR as described previously^15^.

### Ethics statement

The care and use of the animals were approved by and in agreement with the Yale Animal Resource Center and Institutional Animal Care and Use Committee (#2018-21098) according to standards in the *Animal Welfare Act*.

### Viral stocks

MNoV^CW3^ (Gen bank accession EF014462.1), MNoV^CR6^ (accession JQ237823), and MNoV^MNV3^ (accession JQ658375.1) were generated from infectious molecular clones. MNoV^CW3^ is a plaque derived isolate of MNV-1^20^. MNoV^F514I^ was generated by site-directed mutagenesis of the parental MNoV^CR6^ plasmid. Plasmids containing infectious molecular clones were transfected into 293T cells (ATCC) to generate a P0 stock as previously described^15, 20, 51^. The P0 virus was passaged in BV2 cells (gift of H.W. Virgin) to create a P1 stock that was used to inoculate BV2 cells at a 0.05 multiplicity of infection (MOI) for 36 hours to generate a working P2 stock. MNoV^WU23^ (accession EU004668), MNoV^CR3^ (accession EU004676.1), MNoV^CR7^ (accession EU004677), MNoV^S99^ (accession EF531291) were passaged RAW 264.7 cells (ATCC SC-6003) up to six times. RAW cell-derived virus was then expanded one time in BV2 cells as described above. Infected BV2 cell cultures were freeze/thawed, cell debris was pelleted at 1200g for 5 minutes (min), supernatant was filtered through a 0.22 µm filter and concentrated through a 100,000 MWCO Amicon Ultra filter. Virus stocks were aliquoted, tittered three independent times in duplicate, and stored at −80°C until use^15^. Sequencing of the MNoV capsid was performed by PCR amplification from viral cDNA using primers 5’-AACAACTTCACGGTCCAGTCGG3’ and 5’-GCTTGAAAGAGTTGGCTTGGAGC-3’ followed by Sanger sequencing by GENWIZ (South Plainfield, NJ). The P2 stock of MNoV^F514I^ was sequence confirmed by Sanger sequencing. MNoV VP1 alignment was performed with ESpript 3.0 software.

### Cell line culture

BV2 cells, 293T cells, and HeLa cells (ATCC) were maintained in Dulbecco’s modified eagle media (DMEM; Gibco, Gaithersburg, MD) supplemented with 10% fetal bovine serum (FBS; VWR, Radnor, PA), 1% pen/strep (Gibco), and 1% HEPES (Gibco). CD300lf-deficient BV2 cells (clone 1B6, BV2ΔCD300lf) were generated by CRISPR/Cas9^15^. BV2ΔCD300lf cells were complemented with transgenic CD300lf as described previously^15^. HIE cultures (Baylor College of Medicine) were maintained as previously described^8^. BL6/J bone marrow progenitors were isolated from CD300lf^−/−^ or CD300lf^+/−^ mice as previously described^15, 52^. Progenitors were differentiated into BMDMs by plating 10^6^ cells into a 10 cm non-tissue culture treated dish with BMDM media (DMEM, 10% fetal bovine serum, 10% CMG14 conditioned media, 1 mM sodium pyruvate, 2 mM L-glutamine, 1% pen/strep) and incubated for seven days at 37°C and 5% CO_2_^53^. BMDM differentiation was confirmed by flow cytometry staining for F4/80.

### *In vitro* MNoV infections

BV2-WT and BV2ΔCD300lf cells (gift of H.W. Virgin) were seeded at 20,000 cells per well in 96-well plates and infected with diverse MNoV strains at a MOI of 5. After 48 hours, 25µL of CellTiter-Glo (Promega, Madison, WI) was added to each well and luminescence was detected on a Synergy luminometer (BioTek). Experiments were performed in duplicate in at least three independent experiments. BMDMs were infected for 24 hours with MNoV^CW3^ at a MOI 5 for flow cytometry experiments and a MOI 0.05 for plaque assays. Human enteroids were inoculated with HNoV GII.4 from stool filtrate as described previously^8^. Tranfected 293T cells were challenged with HNoV GII.6 from stool filtrate as described below. The HNoV GII.6 stool filtrate was determined to be infectious using BJAB cells as described previously^9^.

### *In vitro* HNoV infections in HIEs

Five-day differentiated jejunal HIE monolayers in 96-well plates were inoculated with HNoV GII.4_Sydney_2012 (2.5×10^5^ genome copies/well) from stool filtrate as described previously^8^. Experiments were conducted in triplicate in three independent experiments. Serial dilutions of polyclonal antibody against huCD300LF (R&D Systems; AF2774) or recombinant human CD300LF/LMIR3 Fc chimera protein (R&D Systems; 2774-LM-050) samples were carried out in CMGF(-) medium containing 500 μM glycochenodeoxycholic acid (GCDCA). An isotype control (Recombinant Human IgG1 Fc, CF; R&D Systems; 110-HG-100) and recombinant rotavirus NSP4 protein were used as controls respectively. GII.4_Sydney_2012 (2.5×10^5^ genome copies) were mixed with an equal volume of media or dilutions of each serum or protein sample at 37°C for 1 hr, and inoculated onto jejunal HIE monolayers for another 1 hr at 37°C in 5% CO_2_. After incubation, monolayers were washed twice with CMGF(-) media to remove unbound virus and cultured in differentiation media with 500 μM GCDCA for the indicated time points. RNA was extracted from each well using the KingFisher Flex Purification system and MAgMAX-96 Viral RNA Isolation kit. RNA extracted at 1 hpi, was used as a baseline to determine the amount of input virus that remained associated with cells after washing the infected cultures to remove the unbound virus. Replication of virus was determined by RNA levels quantified from samples extracted at 24 hpi.

### Mouse infections

Mice were perorally (PO) inoculated with 25µL of 10^6^ PFU MNoV diluted in D10 (DMEM with 10% FBS). For intraperitoneal (IP) challenge, MNoV was diluted to either 10^7^ or 10^6^ PFU per 200µL in phosphate buffered saline (PBS) and injected into the left lower quadrant of the peritoneal cavity with an insulin syringe. Fecal transmission assay was performed by infecting *Cd300lf^−/−^* or *Cd300lf^+/−^* mice with 25µL PO of 10^6^ PFU MNoV^CR6^. At seven dpi, a freshly isolated fecal pellet was homogenized in 100µL of PBS and 25µL of the fecal slurry was administered PO to *Stat1^−/−^* mice. Seven days after fecal transfer, fecal samples from the *Stat1^−/−^* recipient mice were tested for MNoV genomes via qPCR as described below.

### Viral heat inactivation

Viral heat inactivation was performed by diluting MNoV^CW3^, MNoV^CR6^, and MNoV^F514I^ to 10^6^ PFU per 50µL aliquots in D10 within PCR strip tubes. One aliquot of each virus tube was placed in a thermal cycler and heated at the following temperatures for 30 seconds: 37°, 47°, 52°, 55°, 58°, 61°, 62.5°, 64°, 67°, 70°, and 72°C. Samples were then immediately placed on ice, and 5µL of each sample was applied to 20,000 BV2 cells in 95µL in a 96-well plate. Cell viability was measured by CellTiter-Glo 48 hours post-infection as described above^15^.

### Virus quantification by plaque assay

BV2 cells were seeded in 6-well plates at 2 x10^6^ cells/well. Tissue samples were weighed and homogenized with 1.0 mm silica beads (BioSpec, Bartlesville, OK) in 1 mL of D10 with 1% Pen/Strep, and 1% HEPES^38^. Tissue homogenates were serially diluted ten-fold. Media was aspirated off the BV2 cells and samples were inoculated to each well and gently rocked for 1 hour. Inoculum was removed and 2 mL of overlay media (MEM containing 1% methylcellulose, 10% FBS, 1% GlutaMAX (Gibco), 1% HEPES, and 1% pen/strep) was added to each well. Inoculated plates were incubated for 48 hours at 37°C and 5% CO_2_ prior to plaque visualization with crystal violet (0.2% crystal violet in 20% ethanol) as described previously^37^. Plaque assay from cell culture was performed by freeze/thawing infected samples at 0, 12, and 24 hpi followed by serial dilutions as described above^15, 37^.

### Quantitative PCR

MNoV genome copies in fecal pellets and tissues were determined as previously described^15, 54^. Briefly, viral RNA from fecal pellets was extracted using the Quick-RNA Viral 96 Kits according to manufacturer’s protocol (Zymo Research, Irvine, CA). Tissue RNA extraction was performed using TRIzol (Life Technologies, Carlsbad, CA) and purified using Direct-zol RNA MiniPrep Plus according to manufacturer’s instructions (Zymo Research). A two-step cDNA synthesis with 5µl RNA, random hexamer, and ImProm-II Reverse Transcriptase (Promega) was performed^31^. Then, qPCR analysis was performed in duplicate for each of the samples and standard curves generated using MNoV specific oligonucleotides: Probe: 5’ 6FAM-CGCTTTGGAACAATG-MGBNFQ 3’; Forward primer: 5’ CACGCCACCGATCTGTTCTG 3’; Reverse primer: 5’ GCGCTGCGCCATCACTC 3’. The limit of detection was 10 MNoV genome copies/µL. MNoV genome copies detected in tissues were normalized to the housekeeping gene β-β-actin which was detected using murine β-actin oligonucleotides: Probe: 6-JOEN-CACCAGTTC /ZEN/ GCCATGGATGACGA-IABkFQ 3’; Forward primer: 5’ GCT CCT TCG TTG CCG GTC CA 3’; Reverse primer: 5’ TTG CAC ATG CCG GAG CCG TT 3’. The actin limit of detection for qPCR was 100 copies/µL. Undetectable MNoV genomes were set at 0.0001 relative to actin. For HNoV GII.6 qPCR, the primer pair NKP2F NKP2R and probe RING2-TP were used as described previously^55^. For HNoV GII.4 qPCR, the primer pair COG2R /QNIF2d and probe QNIFS were used^56^. A standard generated using a 10-fold dilution series of a recombinant HuNoV RNA transcript was used to quantitate viral genome equivalents in RNA samples from GII.4 infected HIEs.

### Flow cytometry

BMDMs were harvested after mock-inoculation or infection with MNoV. Cells were pelleted by centrifugation at 200 x g for 5 min and suspended in 250µL of Cytofix/Cytoperm (BD Biosciences) for 20 min at RT. Cells were washed twice with perm/wash buffer (PWB), suspended in staining buffer containing rabbit anti-NS1/2 antibody (1:1000; kind gift of Vernon Ward). The cells were incubated with antibody for 30 min at RT, pelleted and washed twice. The cells were suspended in staining buffer containing donkey anti-rabbit alexa fluor 647 (1:500, Life Technologies #A31573) and incubated for 30 min in the dark. The cells were pelleted and washed twice with PWB buffer. Cells were suspended in FACS buffer. Flow cytometry was performed on a MACSQuant Analyzer 10 (Miltenyi Biotec, Somerville, MA) and analyzed using FlowJo v10 (FlowJo LLC, Ashland, OR).

### Mouse CD300 overexpression

HeLa cells were seeded at 400,000 cells per well in a 6-well plate and grown for 24 hours. Murine CD300lf and CD300ld (pcDNA3.4) were transiently transfected into HeLa cells with Trans-It LT1 (Mirus Bio, Madison, WI) according to manufacturer instructions^15^. MNoV^CW3^, MNoV^CR6^ and MNoV^F514I^ were inoculated in transduced HeLa cells at a MOI of 5 and incubated for 24 hours at 37°C. Cells were harvested using Trypsin-EDTA and resuspended in FACS buffer (PBS containing 10% FBS and 2mM EDTA) and stained with anti-CD300lf PE clone Tx70 (1:100; BioLegend, #132704) or anti-CD300ld PE clone Tx69 (1:100, BioLegend, #139605). Cells were then stained intracellularly with guinea pig anti-NS6/7 (kind gift from Kim Green) and goat anti-guinea pig Alexa Fluor 647 (1:500, Life Technologies, #A21450). Cells were then washed twice with PWB and then resuspended in FACS buffer.

### Neutralization Assay

*Cd300lf^−/−^* or *Cd300lf^+/−^* mice were infected PO with 10^6^ PFU MNoV^CW3^ and terminally bled via cardiac puncture at 14 dpi. Whole blood was collected in an EDTA Microtainer tube (BD) and pelleted at 5800 x g for 10 min. Plasma was removed and stored at 4°C until use. BV2 cells were seeded at 20,000 cells in 50µL per well in a 96-cell plate and incubated at 37°C. Sera were serially diluted in D10 with an initial dilution 1:9 and with five subsequent three-fold dilutions. Then, 10^5^ PFU of MNoV^CW3^ was added to each well and mixtures were gently rocked for 30 min at room temperature (RT) and then added to BV2 cells. After 48 hours, 25µL of CellTiter-Glo was added to each well and luminescence was detected on a luminometer. Experiments were performed in duplicate with at least three independent replicates.

### MNoV-specific ELISA

In a 96-well MaxiSorp plate, 100µL of two-fold serially diluted IgG (starting at 12.5ng/mL) and IgM (starting at 25ng/mL) was used as standard controls and MNoV^CR6^ was used to coat the plate overnight at 4°C. The plate was then washed three times with 300µL wash buffer (0.05% Tween-20 in PBS) and then blocked with 100µL of blocking solution (1% BSA in PBS) for 1 hour at RT. After washing 3 times, 50µL of sample at appropriate dilutions in blocking solution was added to each well and incubated for 2 hours at RT. The plate was further washed three times and 50µL of anti-mouse IgG-HRP (A3673, Sigma-Aldrich) or anti-mouse IgM-HRP (A8786, Sigma-Aldrich) diluted in blocking solution was added to each well, then incubated for 2 hours at RT. Wells were washed three times, and then 40µL of ELISA TMB substrate solution (eBioscience, San Diego, CA) was added to each. Plates were incubated for 20 min, then 20µL of stop solution (2N H_2_SO_4_) added and OD determined at 450 nm and reference wavelength 570 nm.

### ELISA for HuNoV VLPs binding to PGM

Pig gastric mucin (Sigma-Aldrich) was immobilized at 10 µg/ml in PBS on 96-well-plate at 4°C overnight. Wells were washed with PBS-0.05% Tween 20 (PBS-T) three times. After blocking with 5% non-fat milk at RT for 1 hour, plates were washed once with PBS-T. Fc-huCD300 proteins (10 µg/ml) were mixed with purified HuNoV VLPs at indicated concentrations and then added and incubated for 1 hour at RT. The wells were washed three times with PBS-T followed by addition of rabbit anti-VP1 sera diluted in PBS (1:2000). Wells were incubated at RT for 1 hour, followed by washing with PBS-T three times as described previously^44^. Secondary goat anti-rabbit-HRP diluted in PBS were added and incubated for 1 hour. After five washes, TMB substrate was added, and 2M H_2_SO_4_ was applied to stop the reaction. Absorbance at 450 nm was measured.

### Statistical Analysis

All statistical analysis was performed in Prism GraphPad version 8 (San Diego, CA). Error bars represent the standard error of the mean unless otherwise indicated. Mann-Whitney tests were performed for all non-normally distributed data whereas normally distributed data was analyzed using Student’s T-tests. All statistical tests were two-sided. Virus heat inactivation was analyzed using a one-way ANOVA. Survival experiments were analyzed by Kaplan-Meier survival curves. A p-value of <0.05 was considered significant (* p-value<0.05, ** p-value<0.01, *** p-value<0.001, **** p-value<0.0001).

## Acknowledgements

We would like to acknowledge Herbert Virgin, Darren Kreamalmeyer, Stephanie Karst, Adam Huys, and Joan Stayos for helpful discussions, generously providing reagents, and technical support. This work was supported by NIH grants K08 AI128043 (CBW), K22 AI151446 (MTB), R01 AI127552 (MTB), R01 AI139314 (MTB), T32 GM007067 (FCW), K99 DK116666 (RCO), and Wellcome Trust [203268/z/16/z] (RSB). CBW was also supported by a Burroughs Wellcome Fund Career Award for Medical Scientists. MTB was also supported by the Pew Biomedical Scholars Program. EAK was supported by NSF Graduate Research Fellowship DGE-1745038. The funders had no role in study design, data collection and analysis, decision to publish, or preparation of the manuscript. CBW and RCO are inventors on a patent application submitted by Washington University entitled “Receptor for norovirus and uses thereof” (U.S. Provisional Application 62/301,965). All relevant data is included in the manuscript.

## Author contributions

VRG, EAK, FCW, EH, JW, KE, MSS, RBF, LLH, AH, and CBW performed experiments. AOK, CEW, LCL, and RSB provided critical reagents. VRG, SMK, MKE, RCO, MTB, and CBW designed the project. VRG, EAK, FCW, EH, JW, KE, AH, SMK, MKE, RCO, MTB, and CBW analyzed data. VRG, MTB, CBW wrote the paper. All authors read and edited the manuscript.

---

**Figure S1.**
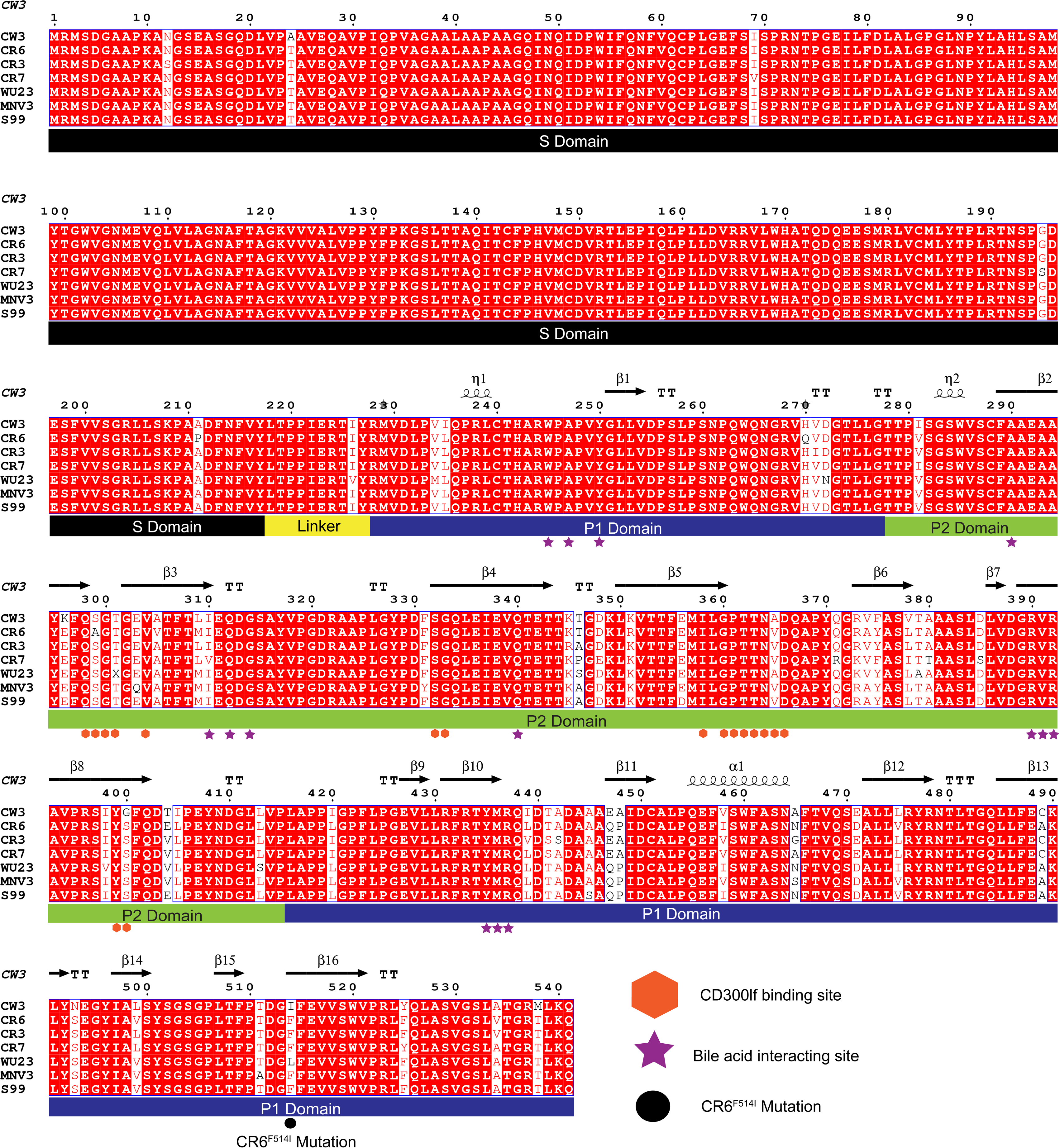
Tree and VP1 alignment showing CD300 interacting sites. VP1 is the major structural protein of MNoV and is comprised of a shell and protruding domain. The complete VP1 sequence of MNoV strains CW3, CR6, CR3, CR7, WU23, MNV3, and S99 were aligned. The VP1 shell domain comprises the core of the virion and is sufficient for virion assembly. The protruding domain mediates binding to CD300lf and bile salts and is comprised of discontinuous P1 and P2 subdomains. The CD300lf and secondary bile acid (GCDCA) binding sites are highlighted as is the F514I mutation which emerged during infection of *Cd300lf^−/−^Stat1^−/−^* mice^13^. Secondary structures labeled as alpha-helices (α), 3_10_-helices (η), and beta-strands (β). The number following the annotation is the numerical order of that secondary structure. Helices are displayed as squiggles and strands are represented by a forward moving arrow under the annotation. TT = strict β-turns and TTT = strict α turns.

